# WhaleLM: Finding Structure and Information in Sperm Whale Vocalizations and Behavior with Machine Learning

**DOI:** 10.1101/2024.10.31.621071

**Authors:** Pratyusha Sharma, Shane Gero, Daniela Rus, Antonio Torralba, Jacob Andreas

## Abstract

Language models (LMs), which are neural sequence predictors trained to model distributions over natural language texts, have come to play a central role in human language technologies like machine translation and information retrieval. They have also contributed to the scientific study of human language itself, enabling progress on long-standing questions about the learnability, optimality, and universality of key features of human languages. Many analogous questions exist in the study of communication between non-human animals—for which, in many cases, we have only a preliminary understanding of signals’ structure and use. Can neural sequence models help us understand these animal communication systems as well? We use these models to characterize the structure and information content of sperm whale vocalizations. Sperm whales (*Physeter macrocephalus*) engage in complex, coordinated behaviours like foraging and navigation in the darkness of the ocean while exchanging sequences of rhythmic clicks known as codas. However, little is known about whether there are any systematic patterns governing coda production, or how codas influence group decision-making and behaviour. To begin to answer these questions, we first train a neural sequence model (a ‘sperm whale language model’) to predict whales’ future vocalizations from their conversational history. By systematically manipulating the information available to this model, and measuring the change in predictive accuracy, we show that sperm whale vocalizations exhibit order dependence, long-range dependencies on up to the past eight codas in an exchange, and predictable turn-taking. Second, we train the sequence model to predict whales’ behaviour from their vocal exchanges, and find that both current behavioural context and future actions are predictable, with accuracies of 72% and 86% respectively, from coda sequences. Our study provides the first evidence that sperm whale vocalizations contain information that could be used to coordinate behaviour. More generally, it offers a framework for using modern machine learning tools for hypothesis generation and to assist in investigating the structure and function of unknown communication systems.

## Introduction

Communication is a key characteristic of intelligence [1–3]. In humans, language allows us to share knowledge, coordinate actions, and establish social structures. Recently, modern language models (LMs)—neural sequence predictors trained to model the probability distribution of natural language text—have advanced our understanding of how efficiency and learnability constraints shape human languages [4–6] as well as scientific understanding of a number of other biological systems [7, 8]. Humans are not the only animals that communicate to coordinate behaviour; non-human organisms produce and perceive communicative signals in very different ways from humans, and many animal communication systems remain incompletely understood. A key factor that has enabled modern LM-guided discovery in other scientific domains outside language is the ability of these models to effectively fit noisy, high-dimensional, sequential data even before human scientists have identified meaningful, interpretable features. Can neural sequence models aid and guide the scientific characterization of animal communication as well?

We use neural sequence models to characterize both the *structure* and *information content* of an animal communication system—specifically, to model communication and behaviour in sperm whales (*Physeter macrocephalus*). Sperm whales exhibit a multi-level social structure [9–11], coordinated group foraging and child-rearing behaviour [12–14], and a complex, socially learned communication system [15–17]. Sperm whale vocalizations consist of sequences of stereotyped, rhythmic click patterns called codas. Several recent studies have characterized codas’ internal structure [18], including with machine learning models [19–21]. But the patterns in which codas are combined into sequences, and their role in coordinating group behaviour, are still not understood.

To obtain first answers to these questions, we train a collection of sperm whale sequence models on several years of recordings from a population of sperm whales in the eastern Caribbean, the EC1 clan. These sequence models receive as input a “conversation history” (a sequence of vocalizations by one or more whales) and attempt to predict either the whales’ future vocalizations, present behavior, or future behaviour. Crucially, we show that these models can be used not just for prediction, but as tools for hypothesis generation and validation. By systematically manipulating the data they are trained on (e.g. by restricting the length of the conversation history they have access to, or hiding specific acoustic features of individual codas), and measuring the impact of these manipulations on predictive power, we can identify which aspects of vocalizations carry information about future vocalizations or behavior.

We first show that vocal exchanges between sperm whales have complex internal structure: coda production exhibits long-range statistical dependencies, and is sensitive to the identity and ordering of the preceding 8 codas (up to 30 seconds in the past)—including not only codas produced by the vocalizing whale, but also those produced by conspecifics. Next, we show that these exchanges contain information about behaviour: sequence models can predict both whales’ present behavioural context and future actions from their vocalizations alone. By inspecting the model’s predictions, we identify a specific, multi-coda motif that is predictive of future diving when made by all whales present in an exchange. These results provide the first evidence that sperm whale vocalizations exhibit long-range structure above the single-coda level, and the first evidence that this structure encodes information about behaviour. More generally, our approach to sequence-model-guided discovery can serve as a precursor to interactive playback experiments by enabling offline hypothesis development; it offers a flexible framework for using the tools of artificial intelligence to understand complex biological systems.

## Method

### Dataset

We study coda exchanges in a manually annotated coda dataset from The Dominica Sperm Whale Project (DSWP). This includes recordings of the Eastern Caribbean clan (EC1) collected between 2014 and 2018 from bio-logging tags (Dtags, [22]) deployed on known individuals off the island of Dominica. This dataset contains manually annotated coda clicks and extracted inter-click intervals comprising 3948 codas [23]. The dataset also contains the accelerometer, gyroscope, and magnetometer readings from the tags. This allows us to compute the position of the tagged whale over time. The EC1 clan has a membership of fewer than 300 individuals [24]. A total of 41 tags were deployed on 25 different individuals in 11 different social units. We conservatively estimate that at least 60 distinct whales are recorded in our dataset. An example sequence of coda exchanges between two whales and the depth profile of the tagged whale is shown in Fig. 1A.

**Fig. 1.**
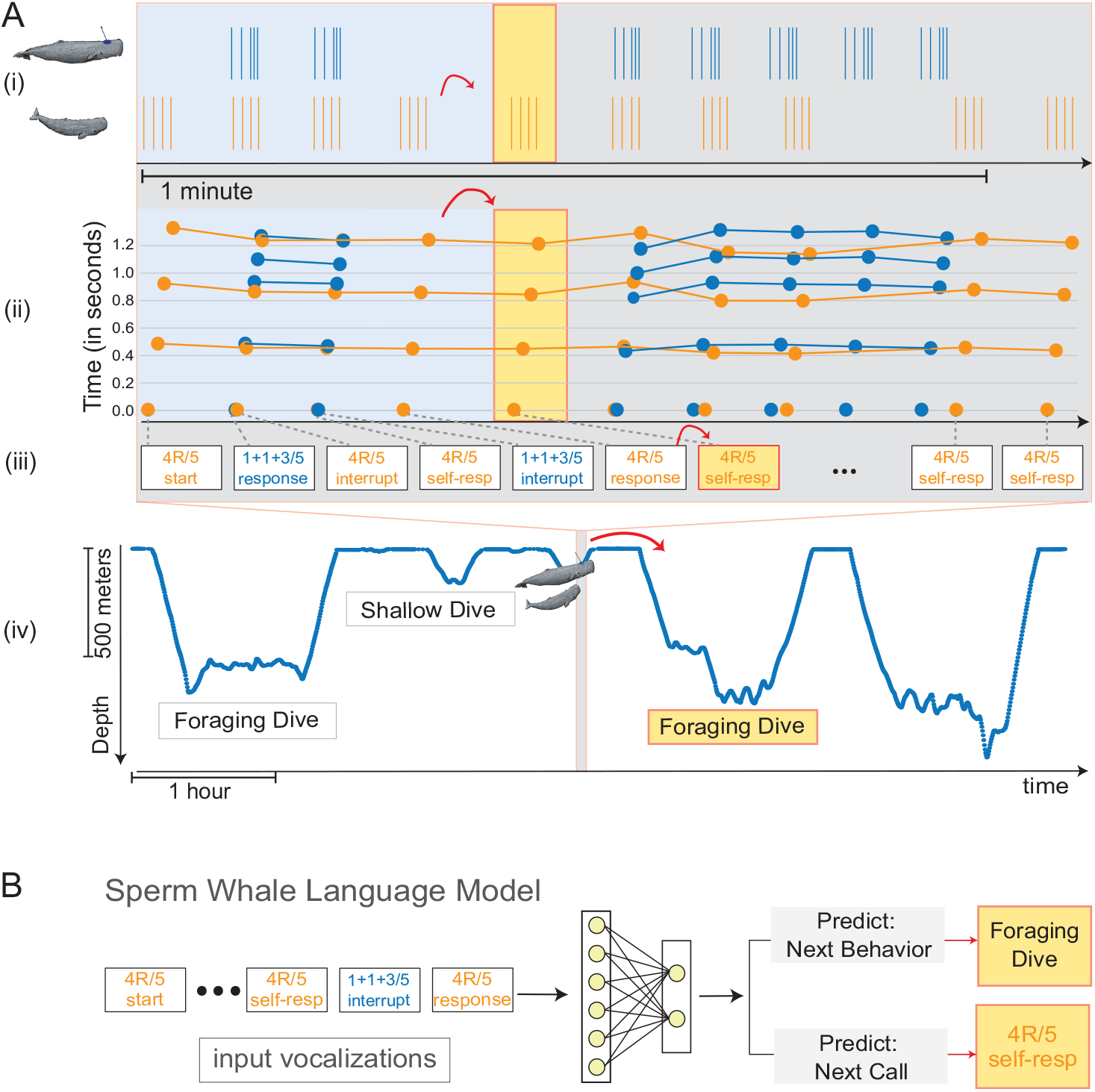
Sperm Whale Language Model. Sperm whales produce sequences of clicks grouped into distinct clusters called codas. **A** shows an (approximately) minute-long interaction between two whales, with the tagged whale’s calls in blue and a conspecific’s calls in orange. (ii) highlights the corresponding exchange plot [23] of the interaction. (iii) depicts this exchange represented as a series of discrete tokens (with codas organized into discrete types following Sharma et al. [23]. (iv) shows the depth profile of the tagged whale over a longer time window. **B** depicts a language model trained on the task of the next coda and behaviour prediction. Like language models trained on human-generated text data, this model is trained autoregressively on sequences of vocalizations like the one depicted in (iii).

### Approach

#### Overview

Using the DSWP dataset, we train a neural sequence model to predict codas and behaviours from preceding coda sequences. We then examine the behaviour of this model to determine what codas and behaviours are predictable and what features of vocalizations support these predictions.

By ‘sequence model’, we refer to any predictive model that takes as input a sequence of discrete observations (which we call ‘tokens’), and places a probability distribution over possible next tokens. For example, a sequence model for a next-word prediction task in English (e.g. ChatGPT [25]) should be able to take the words *the quick brown fox* as input, and judge that the words *jumps over* will occur next with high probability, *sings beautifully* with lower probability, and *purple seventeen* with very low probability. Importantly, sequence models may be trained to perform tasks other than prediction of future tokens, and the input and output spaces need not be the same—a model of the relationship between news reports and stock prices, for example, might take today’s news headlines as input, and place a distribution over yesterday’s *or* tomorrow’s values for some stock market index.

In this paper, we are specifically concerned with sequence models parameterized by deep neural networks, which encode and then predict by first embedding input data in a high-dimensional vector space, then applying alternating linear and non-linear transformations to these token representations mapping them to a distribution over possible outputs. The parameters of the neural network are learned from data as described below. We train two families of neural sequence models, one of which models the distribution over coda sequences, and the other of which models the distribution over behaviours given coda sequences. We begin by formally defining these networks and their training objective, then describe how they can be used to analyze the structure and information content of sperm whale vocalizations.

#### Model training and evaluation

Our dataset (denoted *D*) comprises a sequence of coda exchanges (each denoted *e*_*i*_), each of which in turn comprises a sequence of codas 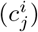, where 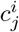 is the *j*th coda in the *i*th exchange. We ‘tokenize’ call sequences by assigning every coda a discrete identifier that captures the four defining coda features (rhythm, tempo, rubato, and ornamentation) previously described by Sharma et al. [23], as well as the time elapsed since the preceding coda in the exchange, and the identity of the vocalizing whale. Each exchange *e*_*i*_ also takes place in a specific behavioural context (e.g. the beginning of a foraging dive or a period of socialization near the surface of the water; see Figure 3). We denote by *b*_*i*_ the behavioral context for the exchange *e*_*i*_. Refer to Section 1.1 in the supplementary for additional details on tokenization.

For each prediction task, we construct an encoder–decoder LSTM [26], a type of recurrent neural network, that maps from a sequence of input codas to a distribution over next codas or behaviour labels. To produce an accurate predictor, we train the network to imitate real coda sequences.

To do so, we first divide the dataset 𝒟 into a training set 𝒟^train^ and a test set 𝒟^test^. When training models for coda prediction tasks, we choose parameters to maximize the log-likelihood 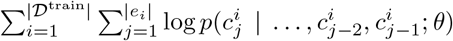, where *p*(*y* | *x*; *θ*) denotes the probability that the LSTM with parameters *θ* assigns to the output *y* given the input *x*. Intuitively, this choice of *θ* encourages the model to assign a high probability to sequences that appeared in the training data and a low probability to all other sequences. When training models for behaviour prediction, we optimize 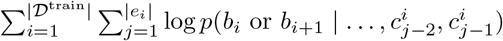, which encourages the model to assign high probability to the true behavioural context of training vocalizations. As described below, our experiments vary both the size of the context window and the features used to distinguish input codas.

As is standard when studying neural sequence models, we evaluate coda-prediction models according to their perplexity exp 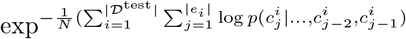, where *N* denotes the total number of codas in 𝒟^test^. Perplexity is simply the exponentiated average log-likelihood per token. Example predictions are shown in Fig. 2E. We evaluate behaviour-prediction models according to their accuracy (whether the behaviour assigned the highest probability matches the ground-truth behaviour in the dataset). Averages for both evaluation metrics are computed over the full DSWP dataset using *k*-fold cross-validation (*k* = 10). Each cross-validation split holds out recordings from a distinct day for evaluation, and trains a sequence model on the remaining days, ensuring that models are evaluated on their ability to extrapolate to novel interactions. Refer to section 1.2 in the supplementary materials for additional training details.

**Fig. 2.**
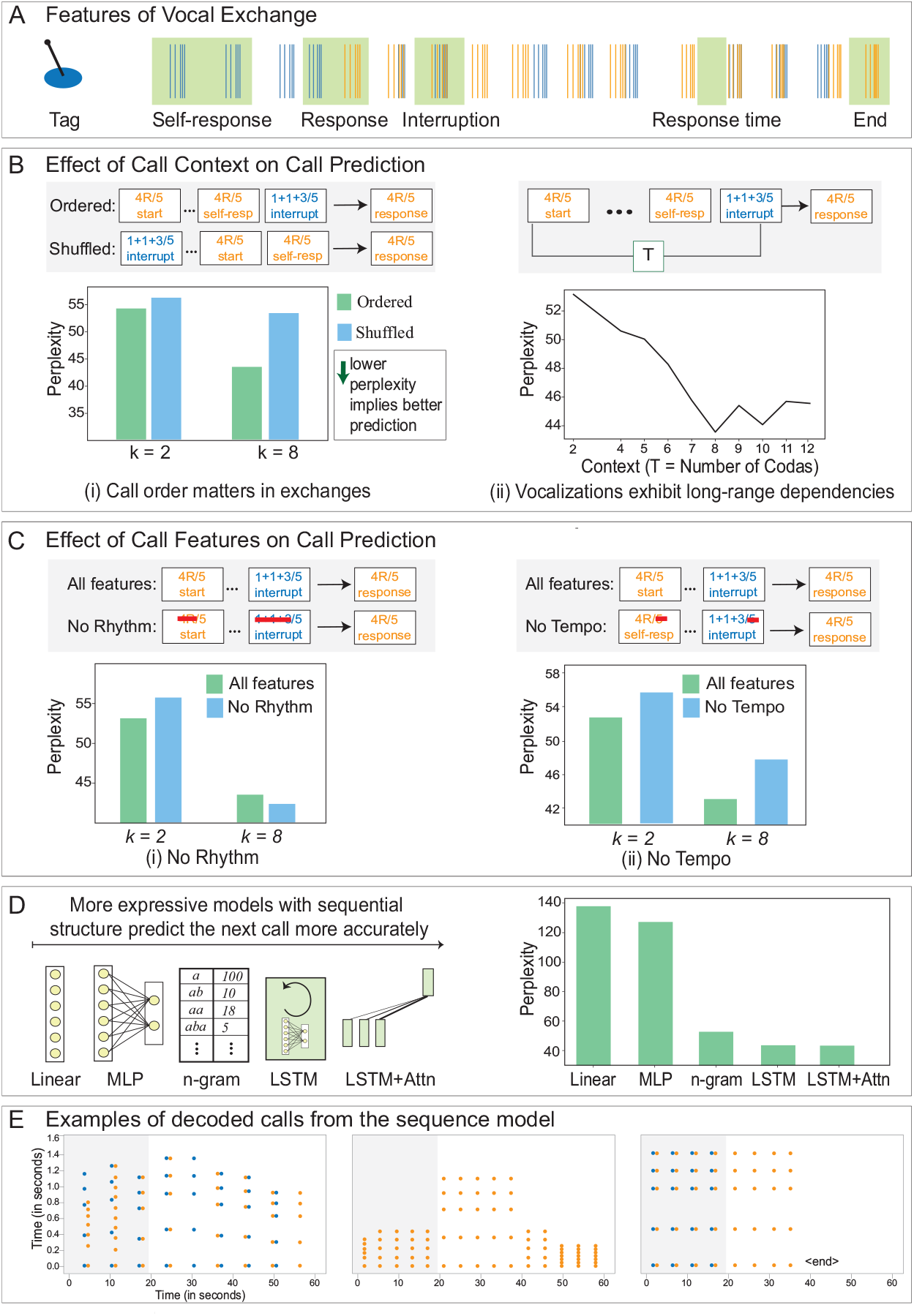
Structure of Sperm Whale Exchanges. We build a sequence model over sperm whale calls to identify what aspects of their call sequence encode information that is predictive of the next call. **A** depicts the sequence of calls exchanged between two whales and various turn-taking and response patterns. In **B**, we analyse the effect of communicative context on call production. We find that coda production is sensitive to the ordering of previous codas (left) and sensitive to a history of up to 8 codas (right). In **C** we evaluate the effect of specific features of the calls on the predictability of the next call in the sequence. We find a considerable decrease in models’ predictive ability when the tempo feature is removed, indicating that the rhythm feature also carries information about future vocalizations. For models with longer input contexts, we observe that omitting the rhythm of the calls has little effect on model performance; however, doing so has a detrimental effect on models with shorter communicative contexts. **D** evaluates the change in model performance with the complexity of the model class. We find that more expressive neural models fit the data distribution better than linear models or (count-based) n-gram models. This shows the existence of long-range dependencies in the communication system that are difficult to model with surface statistics alone. **E** shows example vocalizations generated from the trained model when “prompted” with the sequence of vocalizations shaded in gray.

#### Experimental method

To understand what features of coda sequences contain information about future vocalizations or behaviour, we repeat the training procedure described above while systematically varying the information available to the model. For example, to determine whether the next coda choice is solely influenced by the single preceding coda, we train two models, one of which conditions on a sequence of *n >* 1 input codas *c*_*j*−*n*_, …, *c*_*j*−2_, *c*_*j*−1_, and the other of which conditions on only the most recent coda *c*_*j*−1_. If these two models exhibit similar perplexity on a held-out set, we conclude that longer contexts contain no additional information that is usable for prediction; if the long-context model performs better, we conclude that there is usable information in codas beyond the most recent [27]. This same methodology can be used to evaluate the informativeness of individual *features* of codas; for example, the “rhythm” feature described by Sharma et al. [23]. To do so, we train one baseline model on full coda sequences as above, and one in which each *input* coda’s identity is determined only by its tempo, rubato, and ornamentation features. For both models, we continue to identify *output* codas as before with all four features (to ensure predictions between the two models are directly comparable). If the second, “ablated” model produces less accurate predictions, we may conclude that the rhythm feature contains information useful for prediction.

Importantly, this method for quantifying the informativeness of features is self-supervised: it requires only raw communication (or communication and behaviour) data, without additional labels or interventions from researchers. Below, we use it to identify aspects of sperm whale vocalizations that carry information about future vocalizations, as well as current and future behaviour.

## Results and Discussion

We first use neural sequence models to study the internal structure of coda sequences. To do so, we train next-coda prediction models while removing various sources of information from the input and measuring the effect on predictivity. Results are shown in Fig. 2.

### Vocalizations exhibit long-range dependencies and order-sensitivity

First, we investigate the effect of communicative context by studying how coda *sequencing* influences call production. As motivation, human languages exhibit complex structure in which words and morphemes must be combined and ordered in specific patterns to convey precise meanings: the sentence *The dog in the park was playing* is meaningful, while the sentence *Dog playing park in the the was* is not. Moreover, natural languages exhibit non-local statistical dependencies: if *dog* were replaced by *dogs*, then *was* would need to be replaced by *were* for the sentence to remain grammatical, even though these words are not adjacent to each other in the surface order of the sentence. Sequence-level structure is by no means unique to humans: past studies have shown that songbirds [28, 29], humpback whales [30], and primates [31–33] also produce vocal sequences that exhibit statistical regularities over long distances. In sperm whales, however, the effect of coda order and the distance over which codas exhibit dependencies have not been characterized.

In Fig. 2B(i), we evaluate the informativeness of *call order*. We hold the context window fixed at the past two codas as well as a longer context of eight codas and train LMs on versions of the data in which these input codas arrive in (1) their natural order, or (2) are replaced with a uniformly random permutation of the input. In both cases, models are trained to predict future calls in their natural order. Removing order information from inputs increases perplexity (i.e. decreases predictivity) by up to 22.7%, indicating that ordering information is crucial for predicting future calls (test: Wilcoxon Sign-Ranked Test, sum of the ranks = 55, p-value = 0.001).

In Fig. 2B(ii), we evaluate the informativeness of *context length* by varying the number of preceding codas available to the LM during training and prediction. Short contexts (containing 6 or fewer preceding codas) substantially reduce the predictability of future codas (by up to 20.6%) relative to long contexts (test: Wilcoxon Sign-Ranked Test, sum of the ranks = 54, p-value = 0.002). Together with the ordering information, these results indicate that the patterns governing call production are sensitive to the ordering of a large number of preceding calls. The sperm whale communication system is sensitive to call order and exhibits statistical dependencies across calls separated by as much as 30 seconds (a typical duration for an 8-coda sequence).

### Predicting vocalizations requires complex models and fine-grained coda representations

Having shown that sperm whale call production is sensitive to call history and call order, we next investigate which features within each individual call influence call production, and how expressive sequence models must be to capture this influence.

Sharma et al. [18] previously proposed to analyze codas as a combination of four features termed *rhythm, tempo, rubato, and ornamentation*. In our first experiment, we evaluate which of these features are needed to predict future vocalizations. To do so, we systematically ablate information about these features (one at a time) from the input, while leaving the model’s output space unchanged. For example, to evaluate the role of the rhythm feature, we assign the same input token identifier to all codas that differ only in their rhythm type: for example, 4R/5 (where 4R denotes the rhythm category and 5 denotes the tempo category), 5R/5 and 1+1+3/5 codas are all mapped to the same input token ID. However, output coda IDs are kept unchanged to ensure all models make predictions over the same set of possible output tokens. We then compare the perplexity of this model to models with access to all features. If removing rhythm information from the input increases perplexity, we may conclude that this feature carries information about future vocalizations. Results of this experiment for rhythm and tempo, are shown in Fig. 2C. It can be seen that, when considering a communicative context of only two codas, both features are predictive. Interestingly, with longer contexts, ablating the rhythm feature no longer meaningfully alters predictivity, indicating that it may be somewhat redundant with information conveyed by changes in the other three features over multiple time steps. Corresponding experiments for ornamentation and rubato are provided in Section 1.4 in the supplementary materials.

The preceding experiments have all used a specific recurrent neural sequence model for prediction. Our final coda prediction experiments evaluate the role that the choice of sequence model plays in these findings. To do so, we compare the predictive accuracy of the model in Fig. 2D with four other neural and non-neural sequence models: (a) a **linear** model in which the input sequence is represented by concatenating indicator features for each input coda in order, then mapped directly to a distribution over next codas, (b) a **multi-layer perceptron** which uses the same input representation as the linear model, but passes these inputs through a neural network with an additional hidden layer, (c) an **n-gram** model which predicts next items by counting empirical frequencies of different input coda sequences, and (d) a **LSTM model with attention**, which augments the sequence-to-sequence LSTM model with a single attention head, as in a pointer-generator network [34]. See Section 1.3 in the supplementary materials for implementation details of all models. Results are shown in Fig. 2D: expressive models with explicit sequential structure predict the next call in a sequence more accurately. Surprisingly, n-gram models perform nearly as well as recurrent models, while models based on a (fixed, non-recurrent) input feature representation obtain significantly worse perplexities. The addition of an attention mechanism does not substantially alter predictivity. These results show that vocalizations can be predicted accurately by a range of learned sequence models, but that recurrent neural models enjoy a slight advantage over their classical counterparts.

### Current and future diving behaviour are predictable from vocalizations alone

We next apply neural sequence models to predict not vocalizations, but behaviour. When not floating on the surface of the water, sperm whales in the Eastern Caribbean community alternate between three high-level behavioural states during their active period: conducting deep foraging dives (at depths of over 600 meters), shallow dives (at depths of less than 200 meters), and sleep (during which whales are perpendicular to the surface of the water at depths of less than 100 meters). These behavioural states can be distinguished using accelerometry data that is captured and is aligned to acoustic data captured by the tags. How these behavioral states influence vocalization— whether sperm whales produce different vocalizations in different behavioral states, or whether their vocalizations carry information about *future* behaviors—has been an open question in research on sperm whales for decades.

Using accelerometry data, we automatically annotated the DSWP dataset with the behaviours that accompany vocalizations. These annotations are shown in Fig. 3A: we split foraging dives into their descent and ascent phases, and additionally mark periods of sleep and shallow dives. See Tab. 1 in the supplementary materials for definitions of the behavioural phases and Section 2.1 in the supplementary materials for details of the annotation procedure. Using these annotations, we then train sequence models with the structure described in Fig. 3B(i) and Fig. 3C(i) to perform two prediction tasks. We first predict the *current* behavioural state (i.e. the state of the whale at the moment a particular coda was produced). Because no vocalization occurs during sleep, this involves discriminating between three states: the descent and ascent phases of foraging dives, along with shallow dives. We additionally also predict the *future* behavioural state of the vocalizing whale. We train a model to predict the tagged whale’s next action after the call sequence is produced: whether it will be a deep foraging dive or some other behaviour (e.g. another shallow dive or sleep).

**Fig. 3.**
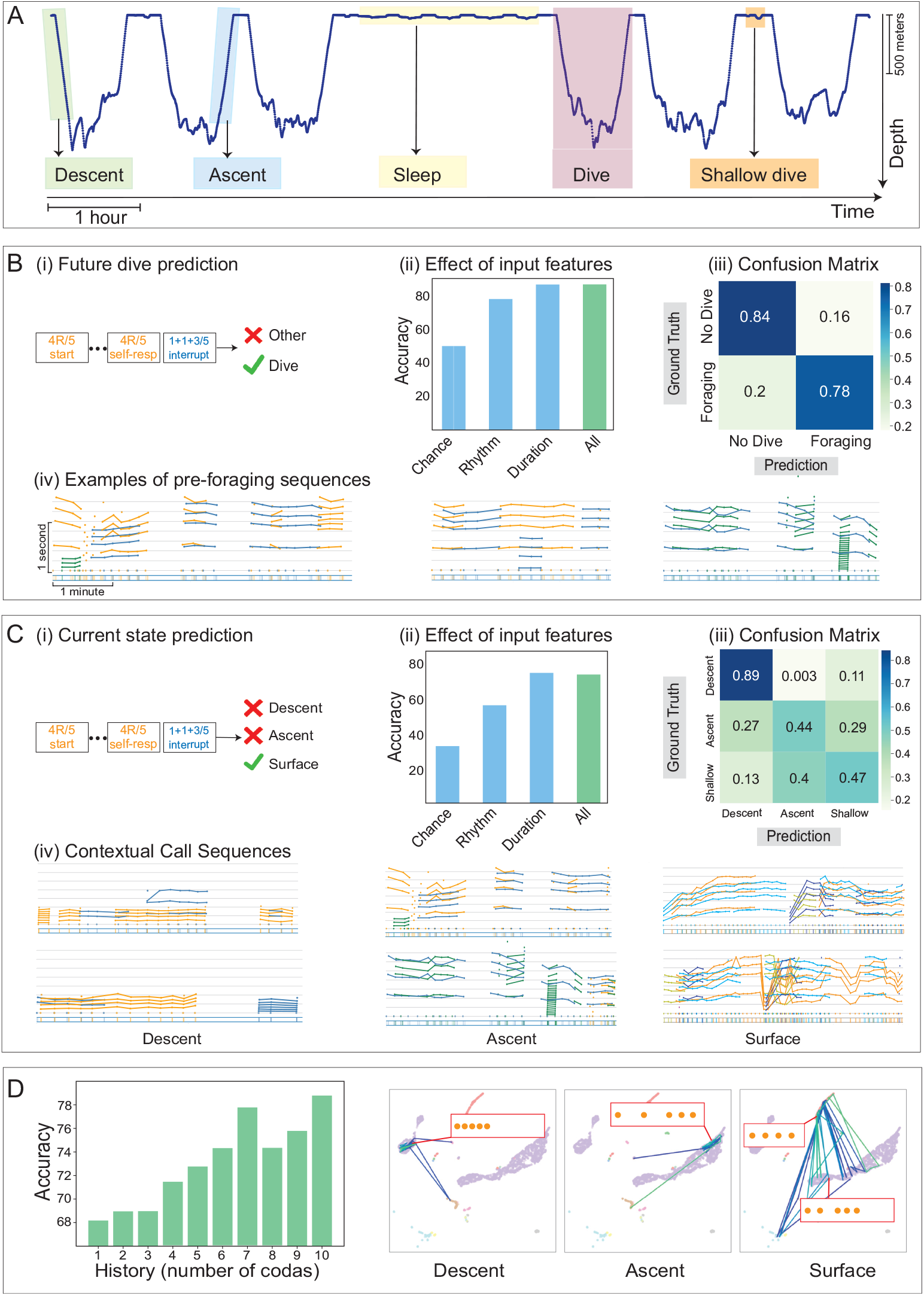
Predicting Behaviour from Vocalizations. **A** shows depth profile of a tagged whale and the corresponding behavioural states of the whale across the period depicted. **B** (i) depicts a neural sequence model trained to predict the future diving behaviour of the whale based on its current sequence of calls. (ii) The model predicts the future behavioural state of the whale correctly 86.4% of the time, significantly better than random-chance baseline of 50%. The sequence of durations of the calls in the sequence is most informative of the next state. (iii) shows the confusion matrix evaluating the model’s performance on the test set over different classes. (iv) Examples of pre-dive codas **C** (i) shows model trained to predict the current state of the whale given a sequence of calls. (ii) The model predicts the current state with an accuracy of 72.8% accuracy, again significantly greater than a chance baseline at 33%. Here too we see that duration information is independently informative about current behavior. (iii) Confusion matrix for the task of current state prediction. (iv) Sample calls for different behavioural states. **D** (Left) Models with a larger input context predict the behavioural state of the whale better. (Right) By embedding codas in two dimensions using t-SNE, and connecting codas produced during the same exchange, we observe characteristic *sequences* of codas associated with different behavioral states, even when some of the constituent codas in these sequences recur across contexts.

Results for both prediction tasks are shown in Fig. 3B(ii, iii) and C(ii, iii). It can be seen that, for both the present and future prediction tasks, a neural sequence model trained to predict behaviour from vocalizations can do so non-trivially, obtaining 72.8% accuracy on the current behaviour prediction task (test: Wilcoxon Sign-Ranked Test, sum of ranks = 54, p-value = 0.002) and 86.4% accuracy on the future behaviour prediction task (test: Wilcoxon Sign-Ranked Test, sum of ranks = 55, p-value = 0.001), compared to a random baseline at 33.3% and 50% for the tasks respectively. It is important to note that we focus on predicting behaviour from vocalizations alone. A sequence model trained to predict future behaviour given only past *behaviour* also obtains nontrivial accuracy. However, the model’s future-behavior predictions are not fully explained by its current-behavior predictions: see Section 2.2 in the supplementary materials for details. Predictability also does not establish a direct causal relationship between vocalization and behaviour, but is a precondition for establishing the influence of communication on behavior.

Inspection of class confusion matrices for these tasks reveals that codas produced during descent are easily distinguished from other behavioural states, while ascent and shallow dive calls are more frequently confused. While [35] shows that the rate of coda production correlates with different sperm whale fluking patterns, our experiments demonstrate that the patterns in which codas are sequenced are distinct across behavioural contexts, and that vocalizations contain information about both present and future behaviour.

### Behaviour prediction is possible from coda sequences, but not isolated codas

As in Fig. 3B(iv) and Fig. 3C(iv), we conclude by evaluating what aspects of whales’ vocalizations are necessary for accurate behaviour prediction. We first evaluate coda-level features by training predictors on rhythm features or tempo features alone. As can be seen in Fig. 3C(ii), tempo features alone suffice to match the accuracy of the full model at both behaviour prediction tasks, while rhythm features alone provide reduced (but still non-trivial) predictive accuracy. This demonstrates that specific combinations of rhythm and tempo are uniquely produced before and during foraging dives.

In the future behaviour prediction task, these results are partially explained by a single coda type that individually predicts future diving behaviour. In Fig. 3B(iv), it can be seen that most pre-dive calls, i.e. calls produced within 15 minutes before the onset of a foraging dive, contain 1+1+3/5 codas (where 1+1+3 denotes the rhythm category and 5 denotes the tempo category), while these are comparatively infrequent in contexts that are not followed by a dive. In fact, considering only the subset of exchanges from Fig. 3B(iv) in which *all* calls (from both the tagged whale and con-specifics) are of the long 1+1+3 type, we find that 67.35% of these exchanges are followed by a dive, while only 19.61% of other exchanges are followed by a dive. While predicting a whale’s current behavioural state improves with a longer input call sequence, the pre-dive call sequences mostly contain a repeated 1+1+3/5 coda (test: Fisher’s exact test (two-sided), odds ratio: 8.94, *p* = 8.2*e*^−7^).

Conversely, for the task of predicting the whales’ current rather than future behaviour, we find that single coda types are not strongly predictive of behavioural context, i.e., ascent, descent and shallow dives: when limiting the number of preceding codas available to the model, as in Fig. 3D(i), accuracy on this task is substantially degraded relative to performance with long input sequences (of seven or more codas). This result indicates distinctive *sequences* of codas discriminate different behavioural contexts from each other. This pattern may again be seen visually: in Fig. 3D(i), we embed all codas from our dataset in two dimensions using t-SNE, then draw lines connecting codas produced sequentially in different behavioural contexts. Each context exhibits a distinctive sequence of coda transitions, even though some individual coda types are produced in multiple contexts. In some cases, these coda sequences are reproduced identically on different days and by different individuals when the same behaviour occurs. If codas are taken to be the atomic units of the communication system, this finding may be interpreted as revealing a kind of “behaviour-dependent syntax” governing coda production (analogous to that observed in house finches [36]).

### Diving is preceded by group signalling

Examining the structure of the pre-dive calls in Fig. 3B(iv), we observe that calls followed by a dive typically involve multiple whales making the same “prototypical” 1+1+3/5 coda. Not only are there signature diving calls, but these calls are 88% of the time produced by multiple whales in the exchange before the diving behaviour is executed. In other words, diving behaviour is distinctively preceded by the production of a single, specific call by multiple members of the unit, a behaviour similar to consensus-formation mechanisms in honeybees [37], birds [38], fish [39] primates [40], and other animals, and not previously described in sperm whales, which may suggest a similar function in sperm whales.

## Concluding Remarks

Machine learning offers promising directions for advancing our understanding the complex communication systems of sperm whales. In this work we have shown that the neural sequence models can identify novel structure within whale vocalizations, predict likely future vocalizations, and in some cases link vocalization to behavior.

A major challenge in studying an animal communication system is simply identifying, from within an enormous hypothesis space, which features of the system are likely to be information-carrying, and how these features relate to behaviour. Our results show that neural sequence models for animal communication, analogous to language models for human languages, can play a key role in meeting these challenges. By training sequence models to predict sperm whales’ vocalizations and behaviour from past vocalization sequences, we have identified informative features and non-local statistical dependencies within sperm whale vocalizations, and provided the first evidence that these vocalizations carry information about whales’ present and future behaviour.

Building on past work that categorizes codas and quantifies production repertoires across social networks and geographic areas (*what* vocalizations sperm whales produce), these new findings place codas in sequential context and link those to behavioural context (*when* and *where* whales vocalize). These results could be extended with the addition of social context describing the identity of vocalizing whales (*who* is producing codas, and for whom). Finally, a key outstanding challenge is grounding vocalization in whales’ own observations of, and goals within, the environment (to understand *why* codas are being produced).

Our findings do not establish what meaning (if any) is derived from a whale’s vocalization by its conspecifics. However, these findings establish, for the first time, two preconditions for understanding if and how sperm whale vocalizations are used to transmit meaning. First, they show that vocalizations are informative: specific vocalizations occur in specific behavioral contexts, making it possible to predict present and future behaviour from vocalizations alone. Second, they show that this information is not localized to individual codas, but distributed across coda sequences, compatible with a higher-level structure analogous to that observed in bonobos [41] and in the syntax of human languages.

The procedure used to obtain these results is general, requiring only sequential representations of communication and behaviour and a large enough dataset to train a sequence model. We thus expect that it could be applied to advance scientific understanding of communication, behaviour, and their relationship across species— especially in ecologically vulnerable populations, like sperm whales, for which tools for *in silico* experimentation can guide hypothesis generation and validation prior to interactive experimentation in the wild.

### Limitations

The machine learning approach described in this paper identifies associational relationships between vocalizations and behaviors. It does not definitively establish the underlying causal structure of the biological system responsible for call production and behaviour coordination, instead providing evidence for “Granger causality” [42]— a statistical association between time series separated in time. Conclusions derived from the predictive power of neural sequence models do not substitute for playback experiments but provide tools to support hypothesis development prior to playback or in populations in which playback cannot be conducted. With more expressive models and larger datasets, it may be possible to extract more elaborate and higher-ordered dependencies in call sequences that improve the predictability of both the next call as well as behaviour.

## Methods

### Data Collection and Coda Annotation

Social units of female and immature sperm whales were located and followed in an area that covered approximately 2000 squared kilometers along the entire western coast of the Island of Dominica (N15.30 W61.40) between 2014 and 2018.

Recordings were made through the deployment of animal-borne sound and movement tags (DTag generation 3, Johnson and Tyack 2003). Tagging was undertaken on an 11-meter rigid-hulled inflatable boat (RHIB). Tags were deployed from a 9-meter, hand-held, carbon fiber pole, and were attached to the whales using four suction cups. DTags record two-channel audio at 120 kHz with a 16-bit resolution, providing a flat (±2 dB) frequency response between 0.4 and 45kHz. Pressure and acceleration were sampled at a rate of 500 Hz with a 16-bit resolution, and were decimated to 25 Hz for analysis. DTag analysis was conducted using custom scripts in Matlab 2015b (The Mathworks, Inc., MA, USA).

Whales, including the tagged whales, were identified through photographs of the trailing edge of their tails [43]. Identifications were used to ensure that only recordings from one of the two sympatric clans (EC1, the Eastern Caribbean Clan) were included in the analysis to control for any differences in repertoires between vocal clans [11].

To define the temporal structure of the codas recorded, absolute inter-click intervals were measured as in [44], using Coda Sorter a custom-written tool (K. Beedholm, Marine Bioacoustics Lab, Aarhus University) in LabView (National Instruments, TX, USA). CodaSorter allows users to play back audio at various speeds and manually mark detected clicks as belonging to a specific coda. Estimates for each click for the angle of arrival, channel delay, centroid frequency, and inter-pulse interval (IPI, the time between the onset of the first pulse and the onset of the next pulse in the multi-pulse structure of sperm whales clicks, [45]) allowed for determining if the codas were produced by tagged whales or non-focal animals; and to ensure that, on days in which multiple tags were deployed, codas recorded by different tags were not double-counted. Photo-identification supported this process by identifying which whales were present and associated with the tagged whales at each surfacing.

The dataset is made up of of 3948 codas which were recorded from animal-borne DTags, which remain in temporal order and are additionally annotated with information of the absolute time in the day of the first click of each of the codas and their associated speaker identities across the exchanges. Rare, long codas were excluded from analysis (greater than 10 clicks, less than 5% of all codas recorded). This dataset was previously used in [23]. Dtag deployments were collected under scientific research permits from the Fisheries Division of the Government of Dominica. The field protocols for approaching, photographing, tagging, and recording sperm whales were approved by either the University Committee on Laboratory Animals of Dalhousie University, Canada; the Animal Welfare and Ethics Committee of the University of St Andrews, Scotland; or Aarhus University, Denmark; and sometimes several or all of these across years.

## Data Availability

The dataset used in the study will be made available upon publication.

## Code Availability

The data was analysed using custom scripts in python. The code will be made available upon publication.

## Acknowledgements

This analysis was funded by Project CETI via grants from Dalio Philanthropies and Ocean X; Sea Grape Foundation; Virgin Unite, Rosamund Zander/Hansjorg Wyss, Chris Anderson/Jacqueline Novogratz through The Audacious Project: a collaborative funding initiative housed at TED.

Fieldwork for The Dominica Sperm Whale Project was supported by through a FNU fellowship for the Danish Council for Independent Research supplemented by a Sapere Aude Research Talent Award (1325-00047A), a Carlsberg Foundation expedition grant (CF14-0789), two Explorer Grants from the National Geographic Society (WW-218R-17 and NGS-64863R-19), a grant from Focused on Nature, and supplementary grants from the Arizona Center for Nature Conservation, Quarters For Conservation all to SG; as well as, small grants from the Dansk Akustisks Selskab, Oticon Foundation, and the Dansk Tennis Fond to Pernille Tønnesen of Aarhus University. Further funding for DSWP fieldwork was provided by a Discovery and Equipment grants from the Natural Sciences and Engineering Research Council of Canada (NSERC) to Hal Whitehead of Dalhousie University and a FNU large frame grant and a Villum Foundation Grant (13273) to Peter Madsen of Aarhus University.

We thank the Chief Fisheries Officers and the Dominica Fisheries Division officers for research permits and their collaboration in data collection; all the crews of R/V Balaena and The DSWP team for data collection, curation, and annotation of the coda dataset; as well as, Mark Johnson and Peter Tyack for in-kind contributions of Dtags during fieldwork in those years; and Dive Dominica, Al Dive, and W.E.T. Dominica for logistical support while in Dominica.

## Author Contributions Statement

P.S., J.A., A.T., D.R. and S.G. conceptualized the study, P.S., J.A., A.T., D.R. developed the methods, S.G. and The DSWP team collected the data, S.G. and The DSWP team annotated the original dataset, P.S., S.G. and The DSWP team curated the data, P.S. conducted the analyses and authored the code provided, P.S., J.A., A.T., D.R., S.G. verified the analytical results, S.G., D.R., A.T., J.A. funded the study, P.S. wrote the manuscript, all authors contributed review and editing to the manuscript and gave final approval for publication.

## Competing Interests Statement

The authors declare no competing interests.

## Notes

### Competing Interest Statement

The authors have declared no competing interest.

### Summary of Updates

The title change was not reflected the first time the manuscript was submitted.

## References

[1] Lieberman, P.: The biology and evolution of language. (1984)

[2] Hauser, M.D., Chomsky, N., Fitch, W.T.: The faculty of language: What is it, who has it, and how did it evolve? Science 298(5598), 1569–1579 (2002)

[3] Jackendoff, R.: Foundations of Language: Brain, Meaning, Grammar, Evolution. Oxford University Press, UK (2002)

[4] Wilcox, E.G., Futrell, R., Levy, R.: Using Computational Models to Test Syntactic Learnability. Linguistic Inquiry, 1–44 (2023)

[5] Kirov, C., Cotterell, R.: Recurrent neural networks in linguistic theory: Revisiting pinker and prince (1988) and the past tense debate. Transactions of the Association for Computational Linguistics 6, 651–665 (2018) 10.1162/tacla00247

[6] Steinert-Threlkeld, S., Szymanik, J.: Learnability and semantic universals. S&P 12, 4–139 (2019)

[7] Jumper, J.M., Evans, R., Pritzel, A., Green, T., Figurnov, M., Ronneberger, O., Tunyasuvunakool, K., Bates, R., Žídek, A., Potapenko, A., Bridgland, A., Meyer, C., Kohl, S.A.A., Ballard, A., Cowie, A., Romera-Paredes, B., Nikolov, S., Jain, R., Adler, J., Back, T., Petersen, S., Reiman, D., Clancy, E., Zielinski, M., Steinegger, M., Pacholska, M., Berghammer, T., Bodenstein, S., Silver, D., Vinyals, O., Senior, A.W., Kavukcuoglu, K., Kohli, P., Hassabis, D.: Highly accurate protein structure prediction with alphafold. Nature 596, 583–589 (2021)

[8] Rives, A., Meier, J., Sercu, T., Goyal, S., Lin, Z., Liu, J., Guo, D., Ott, M., Zitnick, C.L., Ma, J., Fergus, R.: Biological structure and function emerge from scaling unsupervised learning to 250 million protein sequences. Proceedings of the National Academy of Sciences 118(15), 2016239118 (2021) 10.1073/pnas.2016239118 https://www.pnas.org/doi/pdf/10.1073/pnas.2016239118

[9] Whitehead, H.: Sperm whales: social evolution in the ocean. Choice 41(06), 41–3452413452 (2004)

[10] Cantor, M., Shoemaker, L.G., Cabral, R.B., Flores, C.O., Varga, M., Whitehead, H.: Multilevel animal societies can emerge from cultural transmission. Nat. Commun. 6, 8091 (2015)

[11] Gero, S., Bøttcher, A., Whitehead, H., Madsen, P.T.: Socially segregated, sympatric sperm whale clans in the atlantic ocean. R Soc Open Sci 3(6), 160061 (2016)

[12] Marcoux, M., Whitehead, H., Rendell, L.: Sperm whale feeding variation by location, year, social group and clan: evidence from stable isotopes. Mar. Ecol. Prog. Ser. 333, 309–314 (2007)

[13] Cantor, M., Whitehead, H.: How does social behaviour differ among sperm whale clans? Mar. Mamm. Sci. 31(4), 1275–1290 (2015)

[14] Whitehead, H., Rendell, L.: Movements, habitat use and feeding success of cultural clans of South Pacific sperm whales. J. Anim. Ecol. 73(1), 190–196 (2004)

[15] Watkins, W.A.: Sperm whale codas. J. Acoust. Soc. Am. 62(6), 1485 (1977)

[16] Weilgart, L., Whitehead, H.: Coda communication by sperm whales (physeter macrocephalus) off the galápagos islands. Canadian Journal of Zoology 71(4), 744–752 (1993) 10.1139/z93-098 10.1139/z93-098

[17] Rendell, L.E., Whitehead, H.: Vocal clans in sperm whales (Physeter macrocephalus). Proc. Biol. Sci. 270(1512), 225–231 (2003)

[18] Sharma, P., Gero, S., Payne, R., Gruber, D.F., Rus, D., Torralba, A., Andreas, J.: Contextual and combinatorial structure in sperm whale vocalisations. Nat. Commun. 15 (2024)

[19] Leitao, A., Lucas, M., Poetto, S., Hersh, T.A., Gero, S., Gruber, D.F., Bronstein, M., Petri, G.: Evidence of social learning across symbolic cultural barriers in sperm whales (2024)

[20] Beguş, G., Leban, A., Gero, S.: Approaching an unknown communication system by latent space exploration and causal inference. arXiv preprint 2303.10931 (2023)

[21] Goldwasser, S., Gruber, D., Kalai, A.T., Paradise, O.: A theory of unsupervised translation motivated by understanding animal communication. In: NeurIPS 2023 (2023)

[22] Johnson, M.P., Tyack, P.L.: A digital acoustic recording tag for measuring the response of wild marine mammals to sound. IEEE journal of oceanic engineering 28(1), 3–12 (2003)

[23] Sharma, P., Gero, S., Payne, R., Gruber, D.F., Rus, D., Torralba, A., Andreas, J.: Contextual and combinatorial structure in sperm whale vocalisations. bioRxiv (2023) 10.1101/2023.12.06.570484 https://www.biorxiv.org/content/early/2023/12/08/2023.12.06.570484.full.pdf

[24] Vachon, F., Rendell, L., Gero, S., Whitehead, H.: Abundance estimate of eastern caribbean sperm whales using large scale regional surveys. Marine Mammal Science (2024)

[25] OpenAI: ChatGPT: A Large Language Model. https://chat.openai.com. Accessed: 2024-09-16 (2023)

[26] Hochreiter, S., Schmidhuber, J.: Long short-term memory. Neural Computation 9(8), 1735–1780 (1997) 10.1162/neco.1997.9.8.1735

[27] O’Connor, J., Andreas, J.: What context features can transformer language models use? In: Proceedings of the Annual Meeting of the Association for Computational Linguistics (2021)

[28] Searcy, W.A., Soha, J., Peters, S., Nowicki, S.: Long-distance dependencies in birdsong syntax. Proceedings of the Royal Society B 289(1967), 20212473 (2022)

[29] Morita, T., Koda, H., Okanoya, K., Tachibana, R.O.: Birdsong sequence exhibits long context dependency comparable to human language syntax. bioRxiv (2020)

[30] Allen, J.A., Garland, E.C., Dunlop, R.A., Noad, M.J.: Network analysis reveals underlying syntactic features in a vocally learnt mammalian display, humpback whale song. Proceedings of the Royal Society B 286(1917), 20192014 (2019)

[31] Inoue, Y., Sinun, W., Yosida, S., Okanoya, K.: Note orders suggest phrase-inserting structure in male mueller’s gibbon songs: a case study. acta ethologica 23, 89–102 (2020)

[32] Clarke, E., Reichard, U.H., Zuberbühler, K.: The syntax and meaning of wild gibbon songs. PloS one 1(1), 73 (2006)

[33] Leroux, M., Bosshard, A.B., Chandia, B., Manser, A., Zuberbühler, K., Townsend, S.W.: Chimpanzees combine pant hoots with food calls into larger structures. Animal Behaviour 179, 41–50 (2021)

[34] See, A., Liu, P.J., Manning, C.D.: Get To The Point: Summarization with Pointer-Generator Networks (2017). https://arxiv.org/abs/1704.04368

[35] Whitehead, H., Weilgart, L.S.: Patterns of visually observable behaviour and vocalizations in groups of female sperm whales. Behaviour 118, 275–296 (1991)

[36] Ciaburri, I., Williams, H.: Context-dependent variation of house finch song syntax. Animal Behaviour 147, 33–42 (2019)

[37] Frisch, K.V.: The dance language and orientation of bees. J. Anim. Ecol. 38(2), 460 (1967)

[38] Black, J.M.: Preflight signalling in swans: A mechanism for group cohesion and flock formation. Ethology 79, 143–157 (2010)

[39] Sumpter, D.J.T., Krause, J., James, R., Couzin, I.D., Ward, A.J.W.: Consensus decision making by fish. Current Biology 18, 1773–1777 (2008)

[40] Stewart, K.I.J., Harcourt, A.H.: Gorillas’ vocalizations during rest periods: Signals of impending departure? Behaviour 130, 29–40 (1994)

[41] Clay, Z., Zuberbühler, K.: Bonobos extract meaning from call sequences. PloS one 6, 18786 (2011) 10.1371/journal.pone.0018786

[42] Granger, C.W.J.: Investigating causal relations by econometric models and cross-spectral methods. Econometrica 37(3), 424–438 (1969). Accessed 2024-08-05

[43] Arnbom, T.: Individual photographic identification : a key to the social organization of sperm whales. (1987)

[44] Gero, S., Whitehead, H.: Critical decline of the eastern caribbean sperm whale population. PLoS One 11(10), 0162019 (2016)

[45] Møhl, B., Wahlberg, M., Madsen, P.T., Heerfordt, A., Lund, A.: The monopulsed nature of sperm whale clicks. J. Acoust. Soc. Am. 114(2), 1143–1154 (2003)

