## Supplementary file for "WhaleLM: Finding Structure and Information in Sperm Whale Vocalizations and Behavior with Machine Learning"

#### 1 Sperm Whale Language Model

##### 1.1 Tokenization scheme

In natural language processing, *tokenization* refers to the process of representing a text corpus in terms of a finite collection of atomic text units called *tokens.* To build a sequence model of whale exchanges, we apply an analogous tokenization procedure to represent call sequences in terms of a finite set of atomic elements. Following the phonetic alphabet features defined by [1], we represent each coda using rhythm, tempo, rubato, and ornamentation (see Table 1 for definitions). Past work identifies 18 rhythm types, 5 tempo types, 3 rubato types, a binary ornamentation feature. To accurately model the structure of whale exchanges, our tokenization scheme also includes information about turn-taking behaviours, which account for speaker changes and the timing of calls. There are three types of turn-taking: 1) Self-response: A whale follows up its own call after a pause. 2) Response by another whale: A different whale responds after a pause. 3) Overlapping call: A whale produces a call that overlaps with another's. Each unique combination of rhythm, tempo, rubato, ornament, and turn-taking behaviour is assigned a unique token. Not all possible combinations are realized across different data splits. Any combination not present in the training split but appearing in the test split is mapped to the same `<unk>` (unknown) token.

##### 1.2 Details on cross-validation

To ensure robust performance estimates, we conduct each experiment on 10 different dataset splits. In each split, whale calls from a single day are held out for testing, while recordings from the remaining days form the training and validation set. There is no overlap in days between the training and test recordings in any split. The size of the train-val-test datasets is different for different splits of the dataset. This dataset splitting ensures we measure the model's ability to generalize to exchanges from a new day. Due to variability in exchanges across days, the model's performance varies across different splits.

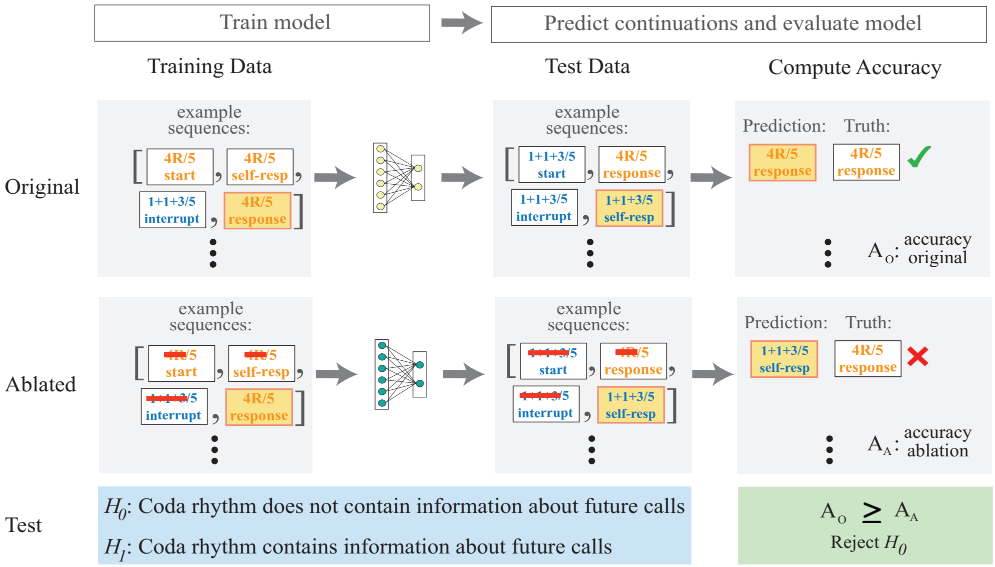


**Fig. 1 Schematic of our method:** Our proposed approach uses sequence models to test hypotheses about the information content and structure of whale calls. Here, we illustrate our method with an example of verifying if rhythm impacts the prediction of future calls. Two models, one with information about rhythm in its input and the other without, are trained and then tested to predict all the call features i.e., the same output. If the model without information of the rubato loses predictive power on the test set, then we reject the null hypothesis "$H_0$: *Coda rhythm does not contain information about future calls*" in favour of the alternate hypothesis.

#### 1.3 Architectural details

To evaluate how expressive sequence models need to be to capture the long-range dependencies and structure of whale calls, we train a collection of models with different inductive biases. Each model is trained on the same input sequence length (sequence length of 6) and optimized for the same objective: predicting the next call. We outline the architectural details of the models below.

##### *n-gram model:*

An n-gram model is a probabilistic language model that predicts the probability of the next item in a sequence based on the previous $n-1$ items. This is done by computing frequency-based estimates of the conditional probability of the next token given the previous $n-1$ tokens on the training set.

The paper uses the implementation of the kenLM repository for training the n-gram models [2] with its default settings. Like with any count-based model, one challenge with an n-gram model is modeling the probability of occurrence of unseen n-grams. To obtain better probability estimates for unseen and less frequent n-grams, Kneser-Ney smoothing is used. Kneser-Ney smoothing starts by discounting from the counts of observed n-grams and redistributes the probability mass to better handle rare and unseen n-grams. Further, the n-gram model is discounted with backoff penalties. Backoff penalties in n-gram models adjust probability estimates when the model has to rely on lower-order n-grams due to the absence of higher-order ones. These penalties help balance the model's reliance on different levels of context, ensuring more accurate and realistic probability estimates across different sequences.

***Linear model:***

A linear model assumes that the output can be expressed as a linear combination of the input features. Here we learn a linear model that outputs the probability distribution over the next token given the previous 6 calls. The number of parameters in this model is a product of the context window times the output.

***Multi-layer perceptron:***

A multi-layer perceptron model contains multiple linear layer layers arranged in a feed-forward fashion with non-linearities between the layers. For this experiment we train a two-layer neural network with a hidden dimension of 64 with a ReLU non-linearity in between.

***LSTM:***

An LSTM (Long Short-Term Memory) is a type of recurrent neural network (RNN) architecture. The LSTM cell contains a more complex unit structure with a specialized gating mechanism that regulates the flow of information in the network, thereby giving it a much more powerful inductive bias to effectively model sequential data. For our experiments, we use an encoder-decoder LSTM, where the encoder LSTM encodes the input sequence into a context vector and the decoder LSTM decodes this vector into an output sequence. We use a bi-directional encoder LSTM cell and a uni-directional decoder LSTM cell, both with 64-dimensional hidden state.

***LSTM with attention:***

An LSTM with Attention is an enhanced version of the Long Short-Term Memory (LSTM) architecture. It incorporates an attention mechanism [] to improve the model's ability to focus on specific parts of the input sequence when generating each element of the output sequence. This architecture is especially useful for tasks when certain parts of the input sequence are more relevant to the output than others. We modify the architecture of the encoder-decoder LSTM to add to this computation.

***Implementational details:***

The parameters of the model were trained with stochastic gradient descent (SGD) using the Adam optimizer with a learning rate of $1e^{-4}$, weight decay of $1e^{-5}$ and batch size of 32. We early-stopped the training of the models based on their performance on a held-out validation set to prevent the models from over-fitting on the small training sets. This usually resulted in the models being trained up to 50 epochs in practice.

#### 1.4 Additional Ablations: Rubato and Ornamentation

Ablating ornamentation and rubato information does not affect the model's ability to predict the next call with statistical significance. This may be partly because ornaments are rare, making up only 4% of the dataset, and the dataset is too small to capture the precise dynamics of changing rubato.

### 2 Behaviour Prediction

#### 2.1 Details on annotating behaviour phases

The different behavioural phases—sleep, shallow dives, and foraging dives—are annotated both automatically and with input from expert humans. Below we outline the procedure used to identify each of the different behavioural phases.

***Foraging dives:***

The start and end points of a whale's foraging dives are moments when the whale starts a sharp descent into the ocean to forage and when the whale first arrives at the ocean's surface by ascending post-foraging. These are automatically detected using the accelerometer and depth data from the DTAG. Foraging dives typically show a steep, uninterrupted descent and ascent profile. A foraging dive is identified when the rate of depth change is nearly constant before and after the start and end points and when the whale reaches a depth of over 500m. This method correctly identifies all the dive start and end points from the collected DTAG data, which are thereafter verified by a human for accuracy.

***Shallow and sleep dives:***

Sleep and shallow dives are identified using accelerometer data and are then annotated and verified by expert human annotators. Examples of the depth and accelerometer data for these dives are shown in Fig. 2. Sleep dives exhibit a distinctive change in the accelerometer reading, indicating the whale's shift from a horizontal position, parallel to the ocean's surface, to a position that is vertical and perpendicular to the ocean's surface. This is highlighted in Fig. 2. In contrast, no such change in the accelerometer reading is observed in other shallow dives. A dive is classified as shallow if the maximum depth of the whale in the course of the dive is less than 100m.

#### 2.2 Behavioural-context baselines for future-behavior prediction.

Our main experiments show that vocalizations contain information about both vocalizing whales' present and future behavior. However, these two prediction targets are correlated with each other: for example, 83% of foraging dives are followed by another foraging dive, rather than a shallow dives. Thus it is possible to obtain non-trivial accuracy at the future-behaviour-prediction task using *only* information about a whale's current state, and not its vocalizations.

In this section, we present an additional analysis showing that these behavioural correlations do not fully explain model accuracy at the future prediction task: that

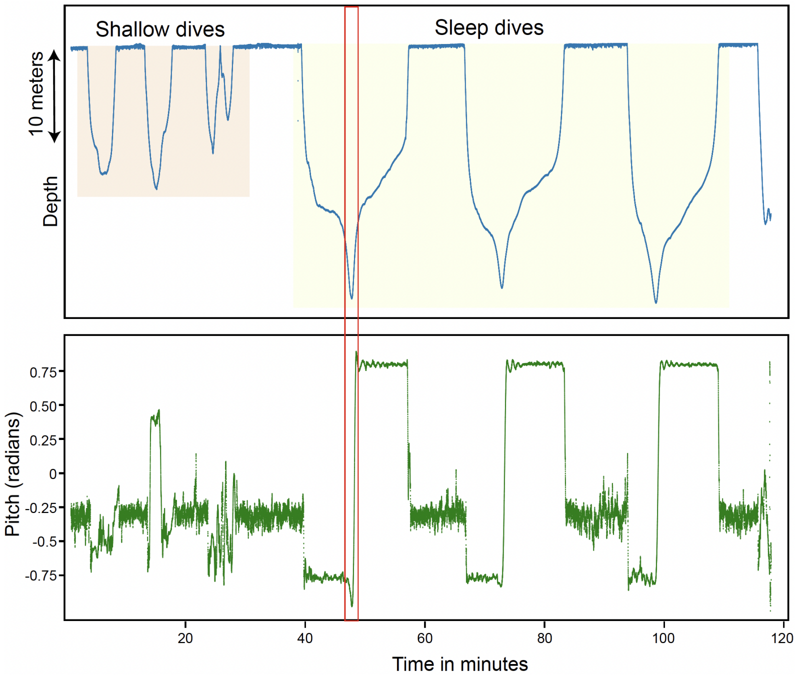


**Fig. 2 Pitch and depth data for shallow and sleep dives:** Shallow and sleep dives are identified using motion data collected using the DTAG. Sperm whale sleep dives have a characteristic change in the depth and pitch data (indicated by the red box) where the whale goes from a position parallel to the surface of the ocean to one where it becomes perpendicular to the ocean.

is, vocalizations contain information about future behaviour *even after accounting for the information they contain about present behaviour.* In particular, we compare the difference in the performance of the future-behaviour-prediction model with a model that predicts the most common next-turn behaviour *conditioned* on a whale's current behavioural state as predicted by the current-behaviour-prediction model. Across the different cross-validation splits, the average difference in the performance between the future-behaviour-prediction model and this suggested baseline model is 21.92% (test: Wilcoxon Sign-Ranked Test, sum of the ranks = 36, p-value = 0.006). This indicates that future-behaviour predictions are not fully explained by correlations between future and current behaviours: some vocalization features are directly predictive of future behavior.

Interestingly, the characteristic "pre-dive" calls identified in the main text are produced not only during the ascent phase but also at the end of social exchanges produced at the surface; however, not all calls produced during ascent follow this "pre-dive" pattern.

| Notation | Description |
|---|---|
| Coda: | A short burst of clicks with varying inter-click intervals generally less than two seconds in duration. |
| Inter Click Interval (ICI): | The time difference between two consecutive clicks within a coda. |
| Coda duration: | The sum of a coda's absolute ICIs. |
| Rhythm type: | The discrete category a coda is assigned to based on its characteristic sequence of standardized ICIs. |
| Tempo type: | The discrete category a coda is assigned to based on its characteristic duration. |
| Exchange / Chorus: | Period of time where codas are made by more than a single whale (as in [3]). |
| Single-Whale Call Sequence: | A sequence of calls made by a given whale where every consecutive pair of calls occur within 8 seconds (twice the average response time) of each other. |
| Turn-taking: | An exchange of codas involving alternating coda production. Also referred to as 'adjacent' codas, these are defined as next-in-sequence codas whose onset occurred within two seconds, but after the termination, of the initial coda (as in [4]). |
| Overlapping Codas: | An exchange of codas such that the next-in-sequence coda's onset occurs after the onset, but before the termination, of the previous coda (as in [4]). |
| Ornament: | "Extra click" appended to the end of a coda in a group of shorter codas. (For further details on the identification criterion, see Ornamentation section in the manuscript.) |
| Rubato: | Gradual variation in duration across adjacent codas made by the same whale within the same rhythm and tempo type. |
| Descent: | The initial period of a foraging dive where there is a steady increase in the depth the whale is located at. This is the period of time starting where the whale is at the surface of the water and makes a plunge to start its foraging dive to the point it reaches a depth at which it can start feeding. |
| Ascent: | The terminal period of a foraging dive where there is a steady decrease in the depth the whale is located at. This is the time period starting where the whale is returning from feeding in deep waters to the point of time it reaches the surface of the water. |
| Social (Socializing on the surface): | The period of time when multiple whales remain at the surface or make shallow dives (¡ 300 meters). |
| Foraging dives: | Deep dives typically involve whales diving to a depth of over four hundred meters. Deep dives almost always have buzzes which are evidence of foraging. |
| Pre-dive calls: | The set of codas made fifteen minutes less before the onet of a foraging dive. |
| behavioural Contexts: | Groups of behaviours exhibited by whales motivated by a set of goals (diving, socializing, pre-dive etc) |
| Context specific calls: | The set of calls prototypically associated with a unique behavioural context. |

**Table 1** Glossary: Definitions of previously used and newly introduced terminology.

| Notation | Description |
|---|---|
| Language: | Any possible set of strings over some (usually finite) alphabet of words. [5] |
| Syntax: | The rules for arranging items (sounds, words, word parts or phrases) into their possible permissible combinations in a language. [5] |
| Likelihood: | Likelihood is a statistical concept that measures how probable a particular set of observations is, given a specific model and its parameters. |
| Neural network: | A neural network is a computational model consisting of interconnected nodes (neurons) organized in layers. It is designed to recognize patterns and learn from data through training by adjusting the connections (weights) between nodes to improve predictions or classifications. |
| Multi-Layer Perceptron: | A multi-layer perceptron (MLP) is a type of neural network consisting of an input layer, one or more hidden layers, and an output layer. Each layer is made up of neurons that use activation functions to process inputs and produce outputs. |
| Neural Sequence Model: | A neural sequence model is a type of machine learning model designed to handle sequential data, where the order of the data points is significant. Sequence models, such as recurrent neural networks (RNNs) and long short-term memory networks (LSTMs), are used to predict the next item in a sequence or to understand dependencies within the sequence. |
| Sequence to sequence models: | A sequence-to-sequence (seq2seq) model is a type of neural network architecture designed to transform one sequence into another. It consists of an encoder that processes the input sequence and a decoder that generates the output sequence. |
| Language Model: | A language model is a type of sequence model typically trained on the next token prediction object. It learns the probabilities of sequences of tokens, enabling it to generate coherent text, autocomplete sentences, or predict the next word in a sentence. |
| LSTM: | A type of recurrent neural network (RNN) architecture designed to effectively capture long-term dependencies in sequential data. |
| n-gram models: | Statistical language models that predict the next item in a sequence based on the preceding $n-1$ items. They represent sequences as contiguous sequences of $n$ items (words or characters) and estimate the probability of each sequence based on the observed frequencies of such sequences in the training data. |
| Perplexity: | Perplexity is a metric used to evaluate the performance of a language model. It is simply the exponentiated average log-likelihood per token. It measures how well the model predicts a sequence, with lower perplexity indicating better performance. |

**Table 2** Glossary 2: Definitions of important linguistics and ML concepts

### Glossary

### References

[1] Sharma, P., Gero, S., Payne, R., Gruber, D.F., Rus, D., Torralba, A., Andreas, J.: Contextual and combinatorial structure in sperm whale vocalisations. Nat. Commun. **15** (2024)

[2] Heafield, K.: KenLM: Faster and smaller language model queries. In: Proceedings of the Sixth Workshop on Statistical Machine Translation, pp. 187–197. Association for Computational Linguistics, Edinburgh, Scotland (2011). https://www.aclweb.org/anthology/W11-2123

[3] Ravignani, A., Bowling, D.L., Fitch, W.T.: Chorusing, synchrony, and the evolutionary functions of rhythm. Front. Psychol. **5**, 1118 (2014)

[4] Schulz, T.M., Whitehead, H., Gero, S., Rendell, L.: Overlapping and matching of codas in vocal interactions between sperm whales: insights into communication function. Anim. Behav. **76**(6), 1977–1988 (2008)

[5] Berwick, R.C., Okanoya, K., Beckers, G.J.L., Bolhuis, J.J.: Songs to syntax: the linguistics of birdsong. Trends in Cognitive Sciences **15**(3), 113–121 (2011) https://doi.org/10.1016/j.tics.2011.01.002
